# Metalation calculator for *E. coli* strain BW25113 grown in M9 minimal media

**DOI:** 10.1101/2025.02.20.639248

**Authors:** Sophie E. Clough, Arthur Glasfeld, Nigel J. Robinson

## Abstract

Here we provide the background data supporting an additional web-based calculator to predict in-cell metalation of proteins within *E. coli* BW25113 grown in M9 media. Intracellular metal availabilities have been estimated from the calibrated responses of DNA-binding, metal-sensing, transcriptional regulators. We have previously tested and validated metalation calculator predictions. They confirm that when proteins are expressed heterologously in *E. coli* in LB medium, they can become mismatched to metal availability and be mismetalated. Mg^2+^GTP-CobW from *Rhodobacter* is predicted to be correctly metalated with cobalt in *E. coli* grown in M9 media, although mismetalated with zinc in LB-grown heterologous cells. Similarly, MncA from *Synechocystis* PCC 6803 is predicted to be correctly metalated with manganese in *E. coli* grown in M9 media, although mismetalated with iron in LB-grown heterologous cells. The M9 metalation calculator is available online and as a spreadsheet for use in optimising metalation in engineering biology. An MncA-refined metalation calculator for idealised cells, reflecting the midpoint of intracellular metal availabilities, is also included and available online.

## Introduction

Metalation calculators have previously been produced for *E. coli* cells grown in LB media aerobically, anaerobically, hyperaerated, exposed to hydrogen peroxide, elevated cobalt, nickel and manganese^1, 2, 3^. The expression of selected genes that are regulated by known DNA binding, metal sensing, transcriptional regulators were previously monitored by qPCR. Boundary conditions describing the highest and lowest levels of transcript abundance for each regulated gene were established. Transcript abundance was related to intracellular metal availability taking advantage of the measured thermodynamic properties of the related metal sensors of *Salmonella enterica* serovar *Typhimurium* (hereafter *Salmonella*). Intracellular metal availabilities were included in on-line calculators that predict which metals will bind to a protein of known metal affinities at the respective metal availabilities when metalation approximates to thermodynamic equilibrium (https://mib-nibb.webspace.durham.ac.uk/metalation-calculators/). The estimated metal availabilities have also been refined based on the occupancies of the metal trap MncA^3, 4^.

Three heterologously expressed proteins (CobW, CbiK and MncA, from *Rhodobacter, Salmonella* and *Synechocystis* PCC 6803) were calculated to be mismetalated within *E. coli* grown in LB media^3^. We hypothesised that this may relate to the rich nature of LB media leading to abnormally high intracellular metal availabilities (expressed as free energies for complex formation in these studies). This manuscript archives the data used to create a metalation calculator based on estimated intracellular metal availabilities within *E. coli* grown in M9 media. The metalation status of the above proteins was re-evaluated under these conditions with implications for biomanufacturing and engineering biology.

## Methods

### Bacterial strain maintenance/growth and reagents

*E. coli* strain BW25113 (hereafter *E. coli*) was grown in liquid growth media and cultures were prepared in acid washed glassware or sterile plasticware to minimise metal contamination. Overnight cultures were inoculated into 400 mL M9 (5x M9 salts, 1 mM MgSO_4_, 0.1 mM CaCl_2_, 1 mM Thiamine-HCl, 20% v/v glucose pH 7.4) at a 1 in 100 dilution and grown at 37 °C to an OD_600 nm_ of 0.2-0.3 then split into 5 mL aliquots in 12 mL capped tubes and grown as indicated. The aerated cultures were incubated with shaking at 200 rpm for 2 h. OD_600 nm_ measurements were made using a Thermo Scientific Multiskan GO spectrophotometer with *n* = 3 independent biological replicates.

### Determination of transcript abundance

Aliquots (1 mL) of culture were added to RNAProtect Bacteria Reagent (Qiagen) (2 mL), RNA isolated a described previously^1^. Briefly, RNA was extracted using an RNeasy Mini Kit (Qiagen) and quantified from *A*_260nm_ followed by DNase I treatment (Fermentas). cDNA was generated using ImProm-II Reverse Transcriptase System (Promega) with control reactions lacking reverse transcriptase prepared in parallel.

Transcript abundance was determined using primers 1 and 2 for *mntS*, 3 and 4 for *fepD*, 5 and 6 for *rcnA*, 7 and 8 for *nikA*, 9 and 10 for *znuA*, 11 and 12 for *zntA*, 13 and 14 for *copA*, 15 and 16 for *rpoD* designed to amplify ∼100 bp (Supplementary Table 1). qPCR was performed in 20 µl reactions containing 5 ng of cDNA, 400 nM of each primer and PowerUP SYBR Green Master Mix (Thermo Fisher Scientific). Three technical replicates of each biological sample were analysed as previously^1^ using a Rotor-Gene Q 2plex (Qiagen; Rotor-Gene-Q Pure Detection Software) with additional control reactions without cDNA templates (qPCR grade water used instead, supplied by Thermo Fisher Scientific) run for each primer pair, in addition to control reactions without reverse transcriptase for the reference gene primer pair (*rpoD*). *C*_q_ values were calculated with LinReg PCR (version 2021.1) after correcting for amplicon efficiency. Change in gene abundance, relative to the control condition (defined as the condition where the minimum transcript abundance was observed for each target gene), was calculated using the 2^−ΔΔ*C*T^ method^5^ using *rpoD* as the reference gene and presented as log_2_(fold change).

### Determination of boundary conditions for the expression of each transcript

Boundary conditions for the calibration of sensor response curves were defined by the minimum and maximum abundance of the regulated transcript which had been previously determined^1^.

Responses of metal sensors (*θ*_D_ for DNA occupancies of metal-dependent de-repressors and co-repressors, *θ*_DM_ for metalated activators) as a function of intracellular available buffered metal concentrations were calculated using sensor metal affinities as described^1, 6^. Transcript abundance was correlated with the response curves to enable estimations of intracellular metal availability expressed as a free energy for complex formation (*ΔG*_M_) in *E. coli* grown aerobically in M9 as described^1, 6^. Using these availabilities, metalation of proteins was predicted *in vivo*, accounting for multiple inter-metal competitions including competition from the intracellular buffer as described by Young and coworkers^7^. Spreadsheets in previous findings from our group perform these calculations for at intracellular metal availabilities in *E. coli* grown aerobically in M9 respectively^3, 7^. The reported values for intracellular availabilities of supplemented metals were thus substituted into Supplementary Data 1.

## Results

Figure 1 shows the fold change in *mntS, fepD, rcnA, nikA, znuA, zntA* and *copA* transcript abundance relative to the level of expression defined previously as the lower boundary condition for the respective promoter^1, 3^. Dashed lines represent maximum expression obtained in elevated cobalt, zinc (for *zntA*) and copper, or depleted iron, zinc (for *znuA*), manganese and nickel (for *nikA*) notably lowest expression of *mntS* transcripts was obtained in cells exposed to manganese and H_2_O_2_. Arrows denote transcript abundance previously obtained under aerobic conditions in LB media^1, 5^. Underlying data (ΔC_q_, ΔΔC_q_ and numerical log_2_ fold changes) are shown in Supplementary Tables 2-4. The fold changes in gene expression shown in Figure 1 were calibrated to DNA occupancies (conditional *θ*_D_ for repressors or *θ*_DM_ for activators) and DNA occupancies related to buffered metal concentration as described previously, to generate Figure 2.

**Figure 1.**
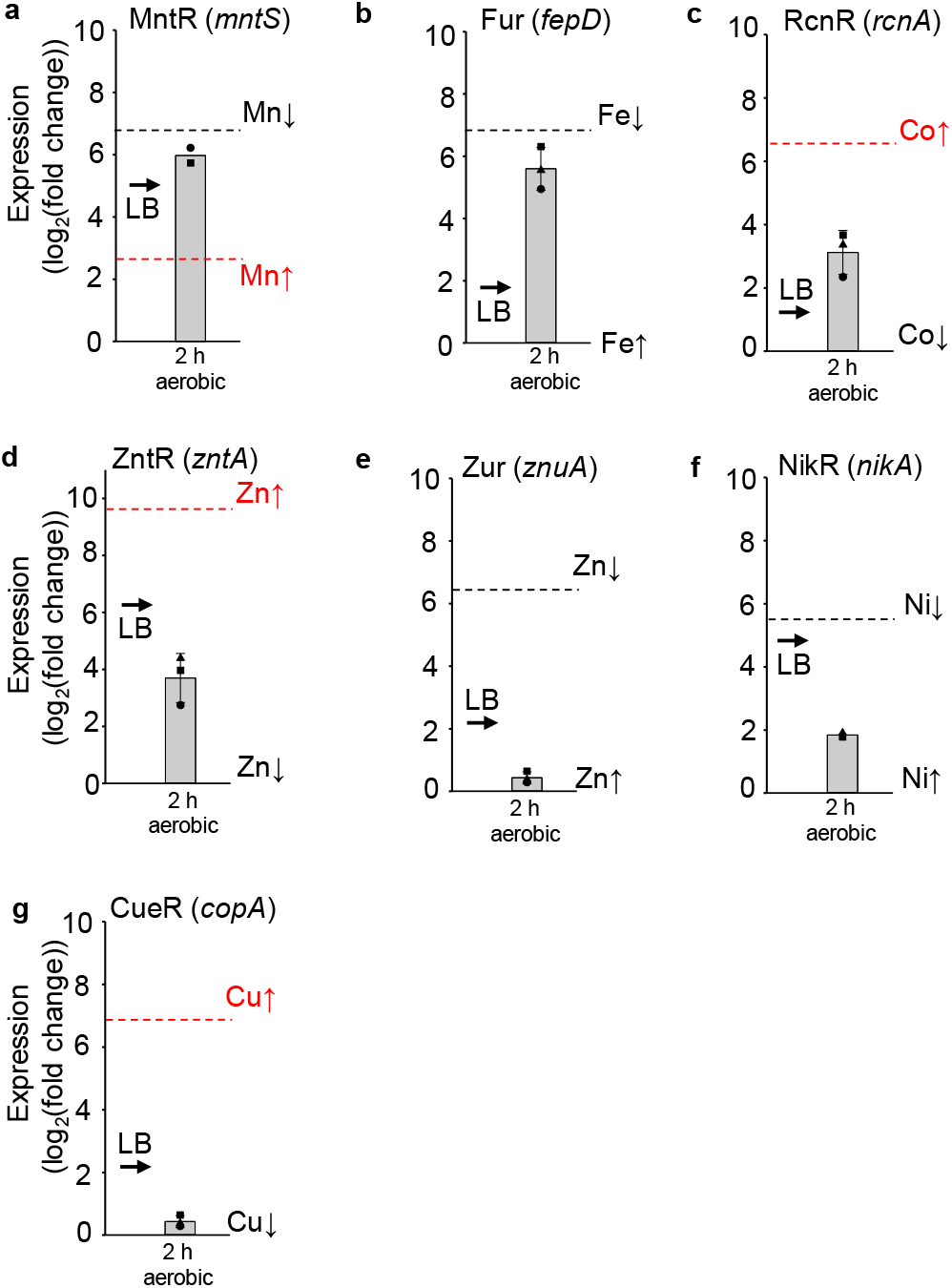
Fold change in metal responsive transcript abundance under hyper-aerated conditions relative to a minimum. As previously for *E. coli* grown in LB media and hyper-aerated^1, 2^, the abundance of seven metal-responsive transcripts were determined in *E. coli* BW25113 grown in M9 medium by qPCR. A. *mntS* regulated by Mn^2+^ sensor MntR. B. *fepD* regulated by Fe^2+^ sensor Fur. C. *rcnA* regulated by Co^2+^ sensor RcnR (noting that RcnR can also response to highly elevated nickel). D. *zntA* regulated by Zn^2+^ sensor ZntR. E. *znuA* regulated by Zn^2+^ sensor Zur. F. *nikA* regulated by Ni^2+^ sensor NikR (aerobic growth). G. *copA* regulated by Cu^+^ sensor CueR.

**Figure 2.**
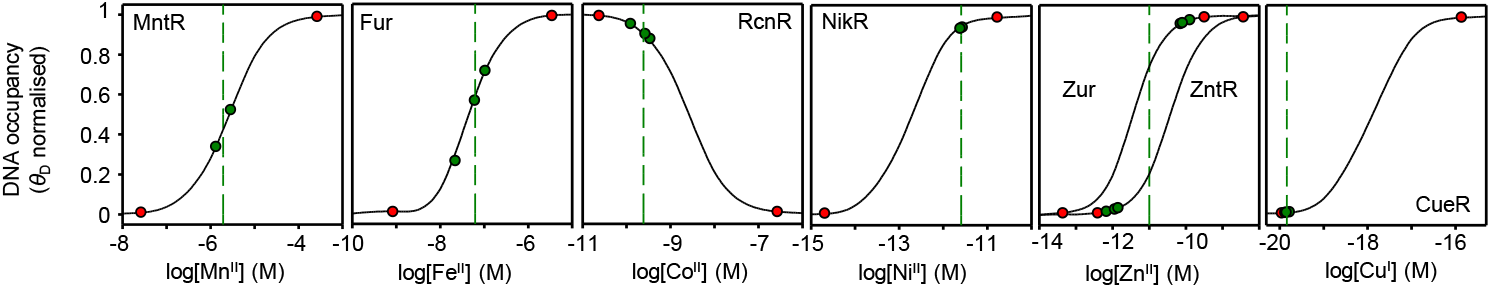
Calibrated responses of metal sensors as a function of intracellular metal-availability on *E. coli* BW25113 grown in M9 media. As previously for *E. coli* grown in LB media and hyper-aerated^1, 2^, The calculated relationship between intracellular metal availability and normalised DNA-occupancy (*θ*_D_), or for activators normalised DNA-occupancy with metalated sensor (*θ*_DM_). The dynamic range within which each sensor responds to changing intracellular metal availability has been defined as *θ*_D_ or *θ*_DM_ of 0.01 to 0.99 representing the boundary conditions for maximum or minimum fold-changes in transcript abundance (red circles), values calculated for *E. coli* cells grown in M9 media (green circles), and mean availability (green dashed lines). For Zn^2+^ two curves represent the responses of *znuA* and *zntA*, here the green dashed line is a midpoint between the means for each sensor. Buffered concentrations and curves were calculated as described by Foster and co-workers^1^. The responses of RcnR to Ni are described separately herein and in^3^.

Buffered concentrations of available metals inside cells grown in M9 media (indicated by the dashed green lines on Fig. 2) were numerically derived from the previously established relationships to *θ*_D_ or *θ*_DM_ for each sensor using MATLAB code (Supplementary Note 3 available in Osman and co-workers^6^), and the values are shown in Table 1.

**Table 1:** Calculated available metal concentrations in cells grown in M9 media^a^.

| Metal | Sensor | M9 cells<br>[Metal]<sub>buffered</sub> (M) |
| --- | --- | --- |
| Mn<sup>2+</sup> | MntR | 2.08E-06 |
| Fe<sup>2+</sup> | Fur | 6.18E-08 |
| Co<sup>2+</sup> | RcnR | 2.42E-10 |
| Zn<sup>2+</sup> | Zur | 9.32E-12 |
|  | ZntR | 1.08E-13 |
|  | Mid-point | 1.0E-12 |
| Ni<sup>2+</sup> | NikR | 2.56E-12 |
| Cu<sup>+</sup> | CueR | 1.46E-20 |

<sup>a</sup>Values were initially calculated to 2 significant figures in Supplementary Data 2 and M9 calculator. Here the finalised numbers have been rounded and the number of decimal places rationalised to those of the standard deviation (shown in parenthesis).

Figure 3 shows metal availabilities inside cells grown in M9 as free energies for forming complexes with proteins (or other types of molecules) that will be 50% metalated at the respective buffered available metal concentrations (Table 1). Standard deviations were calculated based upon the independent experimental determinations which have been averaged in Table 1 (*n* = 3 except for Mn^2+^ and Ni^2+^ where *n* = 2). For comparison, previously determined metal-availabilities at the mid-points of the ranges of each sensor for each metal in idealised cells, are also shown (grey squares in Fig. 3a) and in LB aerobic cells (grey triangles in Fig. 3b). The idealised values shown in Figure 3a incorporate MncA-refined estimations of maximum intracellular nickel availability included in a revised metalation calculator coupled with minimum intracellular nickel availability detected by NikR (Supplementary Data 1). Notably the free energy for complex formation by intracellular available Fe^2+^ is substantially less in cells grown in M9 media compared to LB media. Mn^2+^ are Cu^2+^ are marginally less available while Co^2+^ is marginally more available (note the free energy calculation to produce the y-axis includes a logarithmic term).

**Figure 3.**
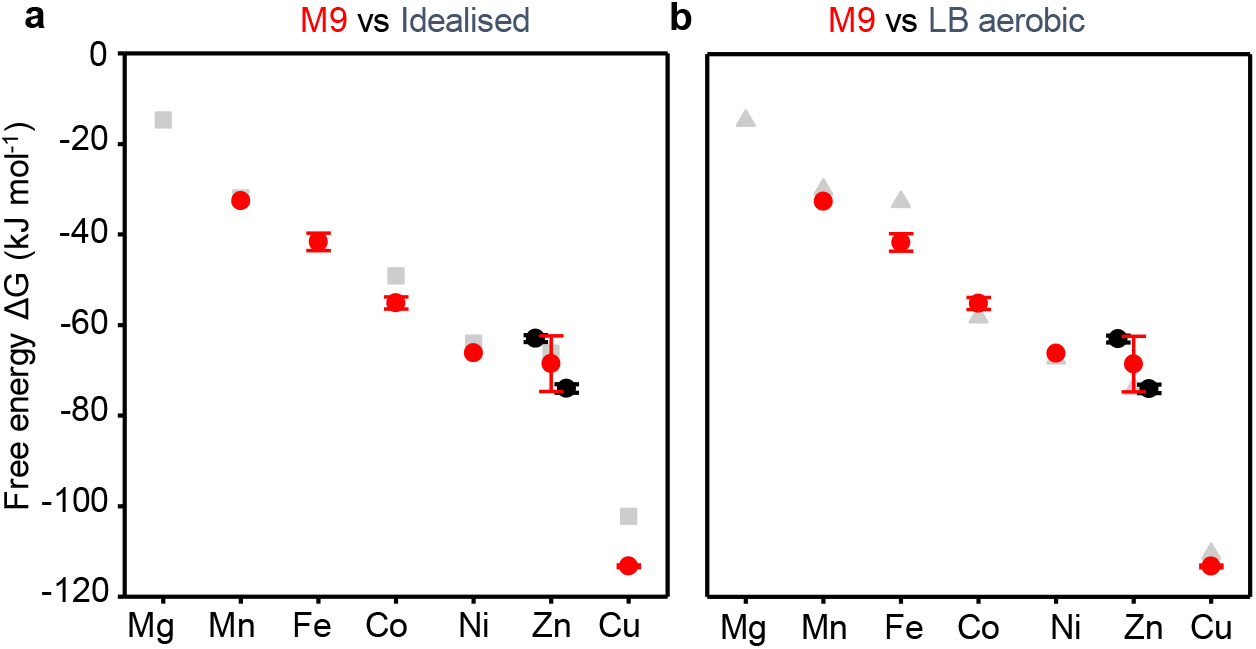
Free energies of available metal in *E. coli* BW25113 grown in M9 media. As previously for *E. coli* grown in LB media and hyper-aerated^1, 2^, Intracellular available free energies for metal binding to molecules that would be 50 % saturated at the available metal concentration (Table 1) in M9 cells (red circles with standard deviation). For Zn^2+^ separate values have been derived based on the response of Zur and ZntR (black circles with standard deviation, Zur = left, ZntR = right). Grey squares in (a) are values for idealised cells reflecting the mid-point of sensor ranges as used in previous versions of calculators and (b) values for aerated cells grown in LB media. The idealised value for Ni^2+^ (6.2 × 10^−12^ M, -64 kJmol^−1^, rather than 1.8 × 10^−13^ M, -72.7 kJmol^−1^ which was based solely on the range for NikR) has been refined using MncA occupancies to define top of range (1.9 × 10^−8^ M, -44.1 kJmol^−1 3^) rather than Ni^2+^RcnR (1.2 × 10^−9^ M, -51.0 kJmol^−1^, *K*_DNA’s_ *and K*_Ni2+_ RcnR determined at pH 7.5), and NikR to define bottom of range (Supplementary Data 1). This refined value is used in Version II of the Metalation Calculator for idealised cells (https://mib-nibb.webspace.durham.ac.uk/metalation-calculators/) and shown in Supplementary Data 1. Intriguingly, using *K*_DNA’s_ *and K*_Ni2+_ RcnR determined at pH 7.0, calculates top of range 6 × 10^− 8^ M, -41.2 kJmol^−1^, remarkably close to the MncA-refined value (Supplementary Fig. 1).

Figure 4 shows the predicted occupancies of five proteins of known metal-binding preferences in cells grown in M9 media using previously described approaches^3^. Figure 5 contrasts the metalation of the cobalt metallochaperone CobW inside cells grown in M9 (calculated using Supplementary Data 2) and LB media, while simultaneously illustrating the outputs of version II of an online metalation calculator which performs the same operations (https://mib-nibb.webspace.durham.ac.uk/metalation-calculators/).

**Figure 4.**
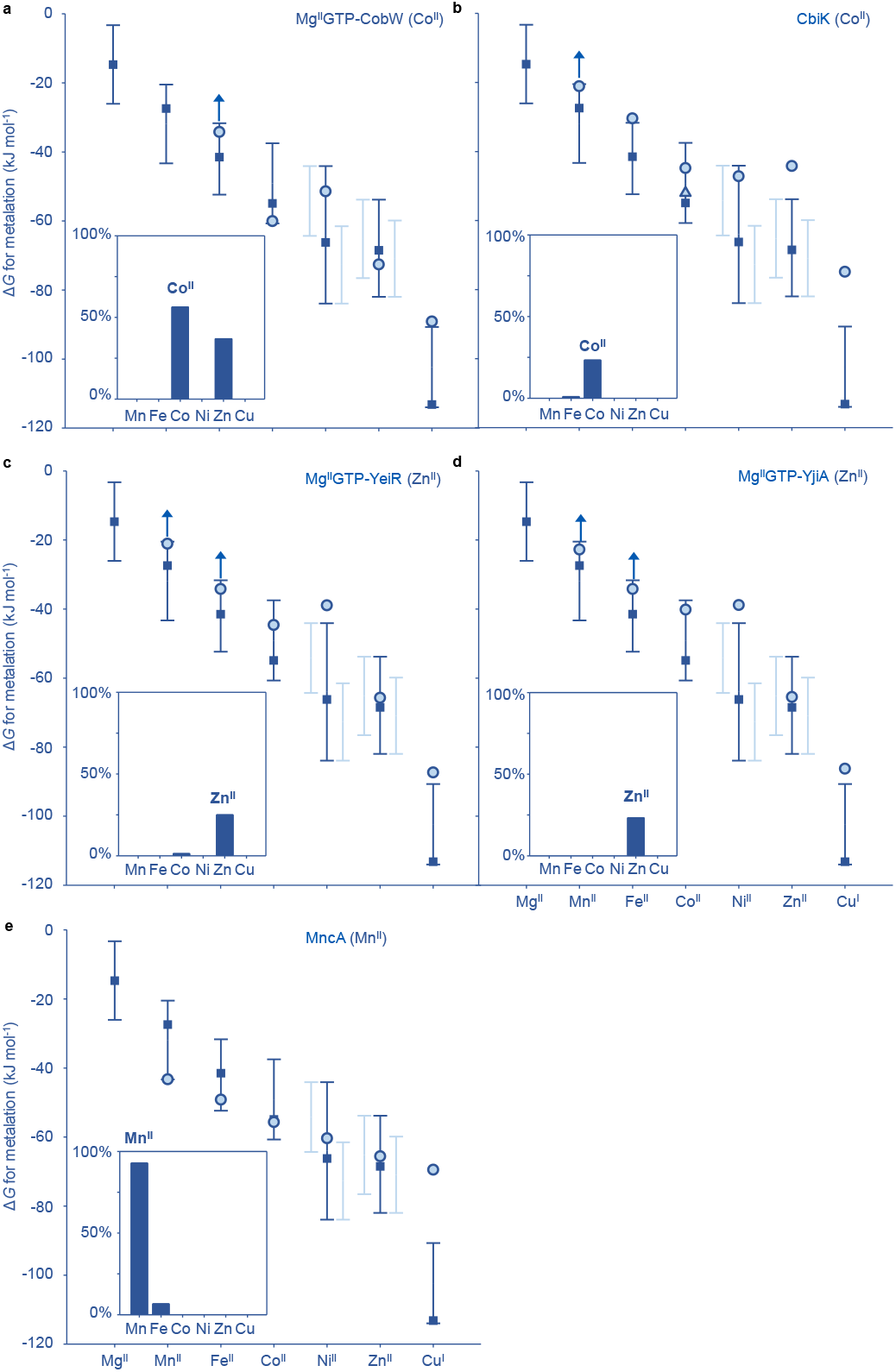
Metal binding preferences of five exemplar proteins compared to metal ion availabilities in M9-grown *E. coli* predict correct metalation under these defined minimal culture conditions. In contrast to *E. coli* grown in LB media^1, 3^, a CobW from *Rhodobacter* bound to Mg^2+^GTP is predicted to be 56% metalated with cobalt. b CbiK, a cobalt chelatase from *Salmonella*, is predicted to be 23% metalated with cobalt using the *K*_m_ values as reported in^8^. c YeiR, a putative zinc chaperone from *Salmonella*, is predicted to be 25% metalated with zinc in its Mg^2+^GTP bound form. d YjiA, a second likely zinc chaperone from *Salmonella* is predicted to be 23% metalated with zinc in its Mg^2+^GTP-bound state. Note that limiting affinities (indicated by small arrows), defined when the protein in question does not compete with the chosen dye in a competition assay, are neglected in the predictions. e MncA, a manganese-dependent oxalate decarboxylase from *Synechocystis* PCC 6803, is predicted to be 93% metalated with manganese.

**Figure 5.**
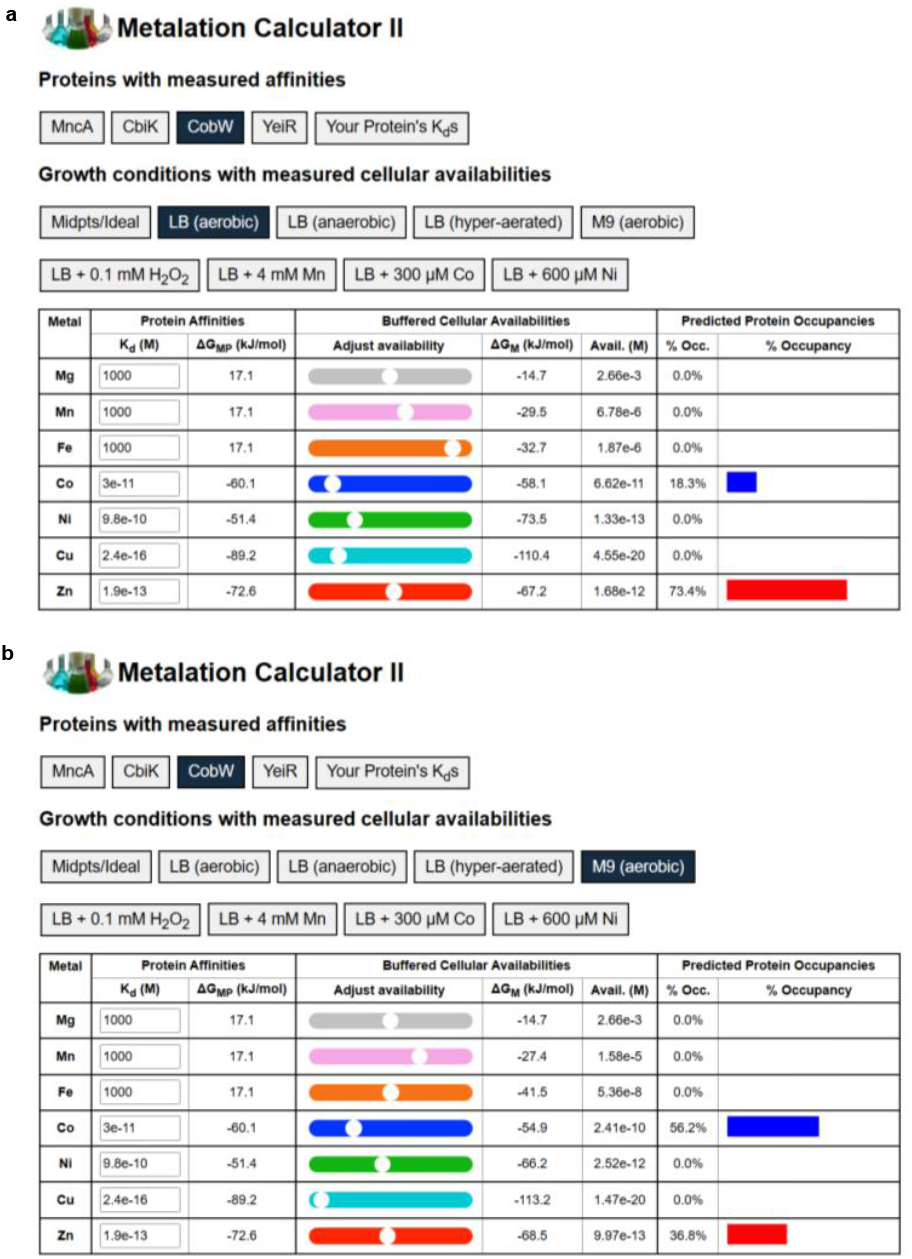
Version II of web-based metalation calculators contrasting metalation of Mg^2+^GTP-CobW in LB versus M9 media. Two screenshots of the metalation calculations available online (https://mib-nibb.webspace.durham.ac.uk/metalation-calculators/). This calculates metal occupancy of a protein or other molecule whose affinities have been entered into the column of editable cells the after selecting “Your Protein’s K_d_s”. Other toggle switches on the top line select specific proteins of known metal-binding preferences such as Mg^2+^GTP-CobW, as selected here (highlighted black). The second and third lines of toggle switches select intracellular metal availabilities such as (a) “LB (aerobic)” (highlighted black) using values from^1^, and (b) “M9 (aerobic)” (highlighted black) using values generated herein. Other availabilities and metal-binding preferences are derived from^1, 2, 3, 6, 7^, plus idealised values for Ni refined as described in Figure 3.

## Discussion

Metal availability in *E. coli* BW25113 cells grown in M9 minimal media, in common with cells grown in LB media, follows the Irving Williams series (Fig. 3, Table 1). Notably, the M9 minimal media provides a source of trace metals even though the recipe contains no specific manganese, iron, cobalt, nickel, zinc or copper salts. It is anticipated that the MgSO_4_ stock provides sources of these ions.

A full complement of metal-binding preferences have previously been determined in vitro for five exemplar proteins: CobW from *Rhodobacter* bound to Mg^II^GTP, CbiK, a cobalt chelatase from *Salmonella*, YeiR, a putative zinc chaperone from *Salmonella* bound to Mg^II^GTP, YjiA, a second likely zinc chaperone from *Salmonella* bound to Mg^II^GTP and MncA, a manganese-dependent oxalate decarboxylase from *Synechocystis* PCC 6803^3, 6, 7^. Relating these preferences to intracellular metal availabilities predicts correct metalation with the cognate metals inside *E. coli* grown in M9 media. This contrasts with previous predictions that the three non-*E. coli* proteins, CobW, CbiK and MncA, will all be mis-metalated (with zinc, iron and iron respectively) when expressed heterologously in *E. coli* grown in LB media^3, 6, 7^. Mismetalation was indirectly evidenced from the CobW-dependent Vitamin B12 synthesis in LB-grown *E. coli* supplemented with cobalt, and directly evidenced by MncA-metal trapping within LB-grown *E. coli*. The current data (Fig. 3-5) highlight the opportunity to rectify mismetalation in engineered cells by adjusting the growth media. This can include supplementation with a cognate metal that is otherwise insufficiently available (such as cobalt for CobW in LB-grown *E. coli*), or depletion of a non-cognate metal that is supra-available such as intracellular iron otherwise outcompeting intracellular manganese for MncA in LB-grown *E. coli* but not in M9-grown *E. coli*. Figure 5 shows how version II of an online metalation calculator (https://mib-nibb.webspace.durham.ac.uk/metalation-calculators/) can assist in identifying which metals need to be manipulated, in which directions, and by how much.

## Supporting information

Supplementary Data 1 (Refined Idealised)

Supplementary Data 3 (RcnR pH 7.0)

## Acknowledgements

This work was supported by Biotechnology and Biological Sciences Research Council awards BB/W015749/1 (N.J.R.), Understanding mis-metalation of native versus heterologously expressed protein, and BB/V006002/1 (N.J.R.), A calculator for metalation inside a cell, along with BB/S009787/1 (N.J.R.) supporting networking in Industrial Biotechnology.

## Data availability

*The data underlying this article are available in the article and in its online supplementary material*.

## Notes

### Competing Interest Statement

The authors have declared no competing interest.

