## Supplementary Information for "Metalation calculator for *E. coli* strain BW25113 grown in M9 minimal media"

**Supplementary Table 1:** Oligonucleotide sequences for qPCR used in this work

| No. | Name | Sequence | Product size (bp) | Reference |
|---|---|---|---|---|
| 1 | *mntS*_F | 5'-GTATGCGCGTGTTTAGTCATTC-3' | 105 | [1] |
| 2 | *mntS*_R | 5'-TATCGGAAGGTTTATCTTGCTG-3' | 105 | [1] |
| 3 | *fepD*_F | 5'-TGCAAACCCTCACCCGAAAC-3' | 111 | [1] |
| 4 | *fepD*_R | 5'-GCGCGGAAGAGTAACCAAACAG-3' | 111 | [1] |
| 5 | *rcnA*_F | 5'-GAACCAGGGCACTCAAAAAC-3' | 108 | [2] |
| 6 | *rcnA*_R | 5'-TGCGGTATGCGAAATAGTTG-3' | 108 | [2] |
| 7 | *nikA*_F | 5'-AACCCGCACCTTTACACGCC-3' | 114 | [1] |
| 8 | *nikA*_R | 5'-AGTCCAGCTTTTTGCCAGCC-3' | 114 | [1] |
| 9 | *znuA*_F | 5'-GTTTGGACTGACACCGCTTG-3' | 111 | [3] |
| 10 | *znuA*_R | 5'-ACGCAGGTTGCTTTTTGCTC-3' | 111 | [3] |
| 11 | *zntA*_F | 5'-CGAAGCACAGGTTGCTGAAC-3' | 114 | [1] |
| 12 | *zntA*_R | 5'-CCGGCAGCGCAAATCAATAC-3' | 114 | [1] |
| 13 | *copA*_F | 5'-GTCACAAACTATCGACCTGACCC-3' | 104 | [1] |
| 14 | *copA*_R | 5'-CATCCGCCTGCTCAACATCC-3' | 104 | [1] |
| 15 | *rpoD*_F | 5'-GTGGCTTGCAGTTCCTTGAC-3' | 108 | [3] |
| 16 | *rpoD*_R | 5'-AGGTTGCGTAGGTGGAGAAC-3' | 108 | [3] |

**Supplementary Table 2:** Δ*C*q values obtained from qPCR analysis of hyperaerated samples with all target primers and *rpoD* (control)[a]

| Target primer | Exposure time (min) | $\Delta C_{q\ target\ gene\text{-}rpoD}$ |
|---|---|---|
| *mntS* | 120 | 0.93 |
| *fepD* | 120 | 2.95 (±0.7) |
| *rcnA* | 120 | 6.37 (±0.7) |
| *nikA* | 120 | 5.34 |
| *znuA* | 120 | 2.54 (±0.4) |
| *zntA* | 120 | 3.56 (±0.9) |
| *copA* | 120 | 2.94 (±0.2) |

[a]Values shown as mean ± standard deviation

**Supplementary Table 3:** ΔΔ*C*q values for conditional response for each promoter. Values shown as mean ± standard deviation[a]

| Target primer | Exposure time (min) | $\Delta\Delta C_{q\ M9\ value\text{-}metallomics\ boundary}$ |
|---|---|---|
| *mntS* | 120 | -5.98 |
| *fepD* | 120 | -5.60 (±0.7) |
| *rcnA* | 120 | -3.12 ( ±0.7) |
| *nikA* | 120 | -1.84 |
| *znuA* | 120 | -1.74 (±0.4) |
| *zntA* | 120 | -3.70 (±0.9) |
| *copA* | 120 | -0.43 (±0.2) |

[a]Values shown as mean ± standard deviation

**Supplementary Table 4:** $Log_2$ fold change values for conditional response for each promoter. Values shown as mean ± standard deviation[a]

| Target primer | Exposure time (min) | $Log_2$ fold change |
|---|---|---|
| *mntS* | 120 | 5.98 |
| *fepD* | 120 | 5.60 (±0.7) |
| *rcnA* | 120 | 3.12 (±0.7) |

| | | |
|---|---|---|
| *nikA* | 120 | 1.84 |
| *znuA* | 120 | 1.78 (±0.4) |
| *zntA* | 120 | 3.70 (±0.9) |
| *copA* | 120 | 0.44 (±0.2) |

[a]Values shown as mean ± standard deviation

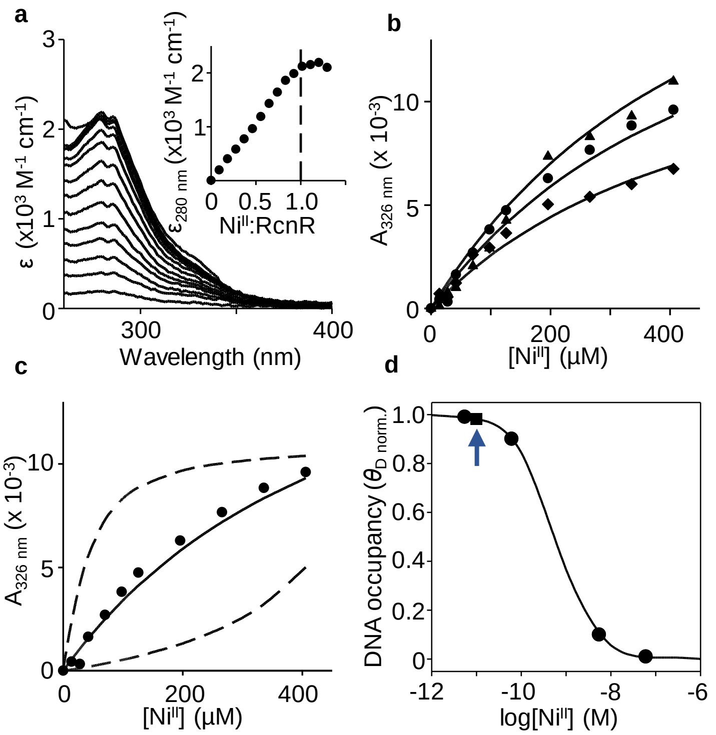


**Supplementary Figure 1. Nickel affinity of RcnR determined at pH 7 to further refine mid-range metal availabilities.** Relatively large residual differences between observed $Ni^{2+}$ occupancy of MncA in nickel-supplemented *E. coli*, and occupancy predicted from the calibrated responses of $Ni^{2+}$-RcnR were previously reported[4]. This could reflect factors not accounted by a thermodynamic equilibrium metalation model, such as kinetic contributions to metalation by $Ni^{2+}$. Alternatively, it could reflect inconsistency in the calibrated Ni responses of RcnR. Notably, values used for $K_{Ni}$ and $K_{DNA}$ of apo-RcnR and $Ni^{2+}$-RcnR were determined at pH 7.5 whereas values determined at pH 7.0 were used for all other sensors. Since the speciation of metalation is a function of relative metal preferences and relative metal availabilities, this disparity might have caused the residual differences. $K_{NiRcnR}$ was therefore determined as described in[4] but using 10 mM HEPES at pH 7.0. **a** Apo-subtracted difference spectra of RcnR (25.8 µM RcnR monomer, ε calculated from total protein) titrated with $Ni^{2+}$ additions from 0 to 35 µM, inset showing peak wavelength confirming 1:1 stoichiometry of $Ni^{2+}$ to RcnR monomer, or 4:1 to $RcnR_4$. **b** Titration with $Ni^{2+}$ to a final concentration of 406 µM in the presence of three concentrations of EGTA and RcnR monomer; 25 µM RcnR (triangle) and 480 µM EGTA; 34 µM RcnR (circle) and 480 µM EGTA; 41 µM RcnR (diamond) and 480 µM EGTA ($n$ = 3 independent experimental trials). **c** Representative $Ni^{2+}$ titration of RcnR in EGTA, solid line representing calculated $K_D$ for $Ni^{2+}$ from simultaneously fitted RcnR-EGTA competitions ($n$ = 3 independent experiments) at varied concentrations of EGTA. Dashed lines, simulations with affinities 10-fold tighter and weaker than calculated $K_D$ ($1.18 \pm 0.07 \times 10^{-10}$ M). **d** $Ni^{2+}$ affinity determined above at pH 7.0 used with previously measured RcnR molecules $cell^{-1}$ and $K_{DNA}$ of $1.5e^{-7}$ M (apo) and $5.9 \times 10^{-6}$ M ($Ni^{2+}$-bound) at pH 7.0[5], to calculate (Supplementary Data 3) relationship between intracellular $Ni^{2+}$ availability and RcnR DNA occupancy ($\theta_D$), as previously done for $Co^{2+}$ (circles show $\theta_D$ 0.99, 0.90, 0.10 and 0.01). Combined mid-point of ranges for $Ni^{2+}$-RcnR and $Ni^{2+}$-NikR also shown ($1.1 \times 10^{-11}$ M, blue arrow).
